# A Recurrent Neural Network Approach for Automated Neural Tracing in Multispectral 3D Images

**DOI:** 10.1101/230441

**Authors:** Yan Yan, Douglas H. Roossien, Benjamin V. Sadis, Jason J. Corso, Dawen Cai

## Abstract

Neuronal morphology reconstruction in fluorescence microscopy 3D images is essential for analyzing neuronal cell type and connectivity. Manual tracing of neurons in these images is time consuming and subjective. Automated tracing is highly desired yet is one of the foremost challenges in computational neuroscience. The multispectral labeling technique, Brainbow utilizes high dimensional spectral information to distinguish intermingled neuronal processes. It is particular interesting to develop new algorithms to include the spectral information into the tracing process. Recently, deep learning approaches achieved state-of-the-art in different computer vision and medical imaging applications. To benefit from the power of deep learning, in this paper, we propose an automated neural tracing approach in multispectral 3D Brainbow images based on recurrent neural net-work. We first adopt VBM4D approach to denoise multispectral 3D images. Then we generate cubes as training samples along the ground truth, manually traced paths. These cubes are the input to the recur-rent neural network. The proposed approach is simple and effective. The approach can be implemented with the deep learning toolbox ‘Keras’ in 100 lines. Finally, to evaluate our approach, we computed the average and standard deviation of DIADEM metric from the ground truth results to our tracing results, and from our tracing results to the ground truth results. Extensive experimental results on the collected dataset demonstrate that the proposed approach performs well in Brainbow labeled mouse brain images.

## Introduction

Analyzing morphology and projection and functional properties of neurons is a key step to understand the connectivity of neurons. As various types of neurons complex connected with each other in a highly intermingled fashion, it is extremely difficult to analyze of neuron morphology in a densely labeled brain. In addition, manual tracing of neurons in 3D images is time consuming and subjective. Therefore, automated neural tracing is one of the foremost challenges in computational neuroscience. To differentiate individual neurons, multispectral labeling, in particular, Brainbow (Cai et al. 2013; Livet et al. 2007) has demonstrated great potential in distinguishing neighboring neurons using random expression of fluorescent protein combinations. With the ability to label each neuron with a distinctive color, it is feasible to reconstruct specific neuronal morphologies and underlying neural circuits Fig. 1. Although attempt has beenmade to segment Brainbow labeled neurons (Sümbül et al. 2016), automated neuronal tracing in Brainbow images remains a challenging task because (i) neurites are nonuniformed branching structures and (ii) multispectral 3D fluorescent Brainbow images normally have high noise level.

**Figure 1.**
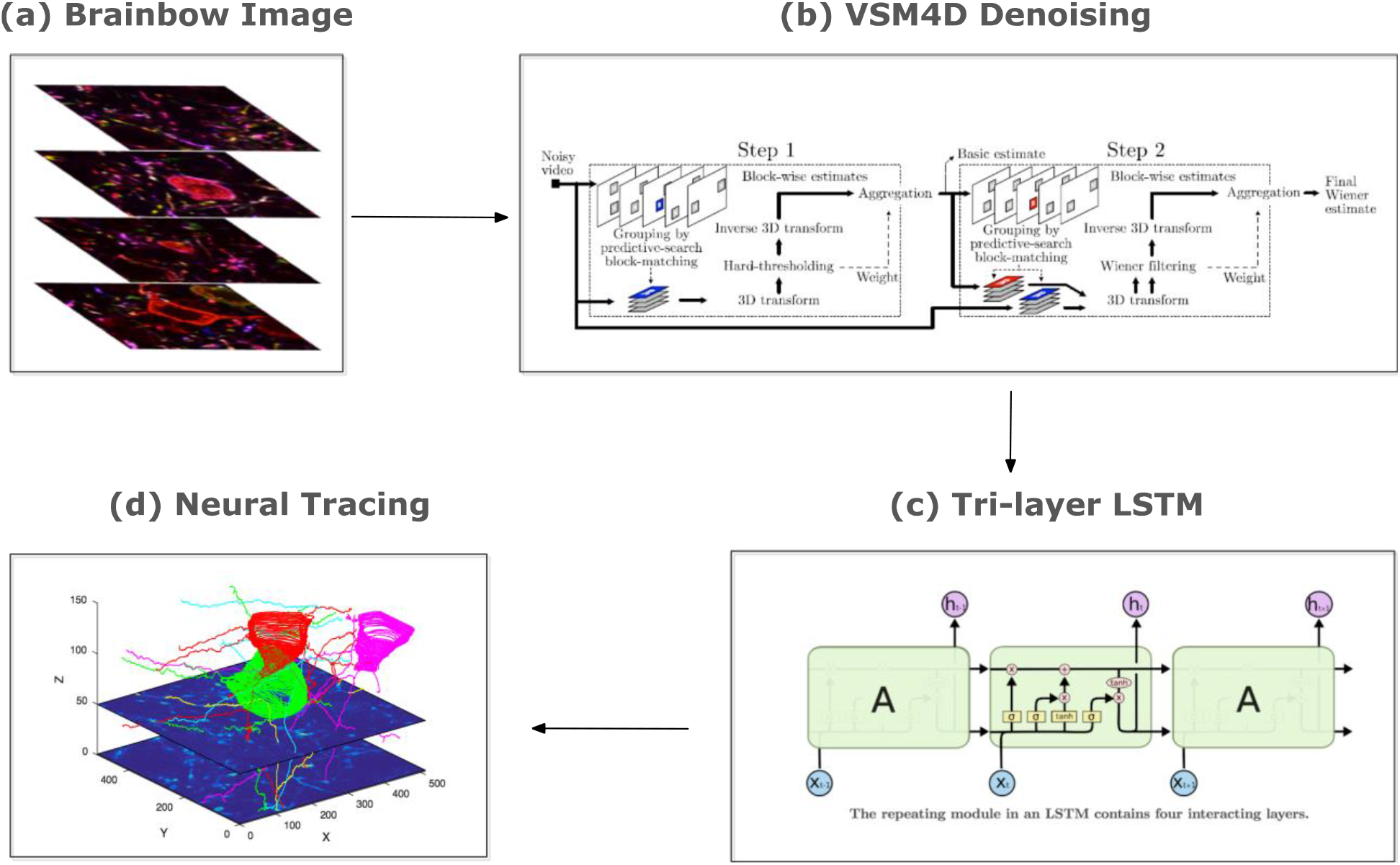
Overview of our proposed frustratingly easy automated neural tracing framework. (a) Input Brainbow Images from the collected dataset. (b) VSM4D Denoising for Brainbow images. (c) Tri-layer LSTM training with cubes. (d) Neural tracing based on learning model. Figure is best viewed in color and under zoom.

In general, neural tracing can be categorized into global processing, local processing and meta-algorithm approaches (Acciai, Soda, and Iannello 2016). Global processing approaches usually process an image as a whole. Wang et al. (Y. Wang et al. 2011) proposed a 3D neuron tracing algorithm based on open-curve active contour, also named as open-curve snake model. Myatt et al. and Longair et al. (Longair, Baker, and Armstrong 2011; Myatt et al. 2012) presented two semi-automatic approaches to manually set both the start and the end points of the neurite. Xiao et al. (Xiao and Peng 2013) proposed an automatic algorithm for neuron tracing based on hierarchical pruning of a gray-scale weighted image distance tree. Local processing approaches explore neural image with relevant local structures. Zhao et al. (Zhao et al. 2011) proposed a local tracing algorithm followed by the shortest path algorithm to optimize the trace. Choromanska et al. (Choromanska, Chang, and Yuste 2012) presented an algorithm that progressively extends the neuronal tree starting from a seed point and analyzing a set of local morphological properties. Bas et al. (Bas and Erdogmus 2011) proposed a method relying on the notion of principal curves. Meta-algorithm approaches do not rely on a particular tracing algorithm, but rather utilize and compare many existing methods in handling large-scale images. They propose a strategy to apply any tracing algorithm designed either for reducing the computational workload or for dealing with image variability. Zhou et al. (Zhou et al. 2016) proposed a fully automatic tracing strategy to gain efficient computation for large-scale images. It first traces the projections on 2D planes, and then reconstructs the 3D neuronal tree using a reverse mapping technique. Chen et al. (H. Chen et al. 2015) developed an automatic tracing framework aiming at overcoming the variability among methods given by the differences in image modality, image parameters or tissue processing protocol.

Nonetheless, most of the existing algorithms are designed to handle monochrome images or omits the spectral information in multispectral images. In addition, although some of above neural tracing approaches are based on machine learning strategies, few of them use deep learning approach which has been successfully adopted in many applications recently, such as image classification (Krizhevsky, Sutskever, and Hinton 2012; W. Wang, Yan, et al. 2016), object detection (Girshick et al. 2014), health question answering (Nie et al. 2015), face and pose analysis (W. Wang, Cui, et al. 2016; Yan et al. 2016), video analysis (J. Chen et al. 2016) and image segmentation (Long, Shelhamer, and Darrell 2015; Yan et al. 2017; Zhang et al. 2013; Zhang, Gao, et al. 2014; Zhang, Song, et al. 2014). Recurrent Neural Network (RNN) is an effective model to process sequential data. RNN can learn complex dynamics by mapping the input sequence to a sequence of hidden variables. By passing the hidden variables recursively to the repeating module in the network, RNN is able to memorize the previous information. Thus, RNN performs well in dealing with sequential data which have dependencies. In the past few years, RNN has been successfully applied to a variety of natural language processing and image processing tasks. Long Short Term Memory (LSTM) networks (Hochreiter and Schmidhuber 1997) are a special kind of RNN, capable of learning long-term dependencies. LSTMs are explicitly designed to avoid the long-term dependency problem. Remembering information for long periods of time is practically their default behavior. They work tremendously well on a large variety of problems, and are now widely used. In this paper, we investigate automated multispectral neural tracing based on LSTM. We extract small cubes along the manually traced path of neural as training samples. For Brainbow 3D images, pixel values from different channels are concatenated as the feature input to LSTM. By capturing the long-term dependencies along neurites, LSTM demonstrates the feasibility of being an effective model for neural tracing. To evaluate our proposed approach, we manually traced a dataset that contain 3D Brainbow labeled neuronal images as the ground truth for the LSTM task. This dataset will be available to the public for future neural tracing research.

In summary, we make two major contributions as following:

- An effective approach based on LSTM is proposed for automated neural tracing in multispectral 3D Images. To the best of our knowledge, we are the first to explore deep learning approach based on RNN for neural tracing.
- A Brainbow 3D image dataset for neural tracing with manual tracing ground truth is provided to the public for future neural tracing research.

## Methods

The overview of our proposed automated neural tracing framework is shown as in Fig.1. This framework is a combination of two key components: a denoising step and a RNN step, respectively. In the image denoising step, we adopt VBM4D denoising strategy (Maggioni et al. 2012; Sümbül et al. 2016) which is considered as state-of-the-art denoising approach for image volumes. In the RNN step, we perform Long short-term memory (LSTM) for the automated neural tracing task. Section 2.1 introduces the image denoising and Section 2.2 presents the RNN.

### Denoising

Being able to collect multispectral information and providing 3D optical sectioning ability in scattering tissues, point scanning confocal microscopy has long been the choice of imaging modality for neuroscience. However, confocal microscopy images are usually noisy due to the inherent characteristics of the detector and contain intensity variations due to tissue scattering especially in depth. The imaging noise and intensity variation can interfere with the biologists’ research by making it difficult to observe low intensity signals and fine detail. Moreover, this limited using methodologies for high-level processing, such as image segmentation and neural tracing.

VBM4D (Maggioni et al. 2012) is a state-of-the-art algorithm for white noise removal in video which evolved from VBM3D (Dabov, Foi, and Egiazarian 2007) by exploiting similarity between 3D spatial-temporal volumes instead of 2D patches and grouped similar volumes together by stacking them along an additional fourth dimension, thus producing a 4D structure. Collaborative filtering is realized by transforming each group through a decorrelating 4D separable transform and then by shrinkage and inverse transformation. Brainbow are 3D images whose depth information (z-axis) surresponses to time information in the video. We performed VBM4D denoiser on each spectral channel of Brainbow images since Brainbow is multispectral image. The results of VBM4D denoising are shown in Fig.2. We can observe that images are smooth and less noisy after VBM4D denosing step. This denoising step greatly improved the tracing accuracy, therefore has been included in our automated pipeline (Fig. 1b).

**Figure 2.**
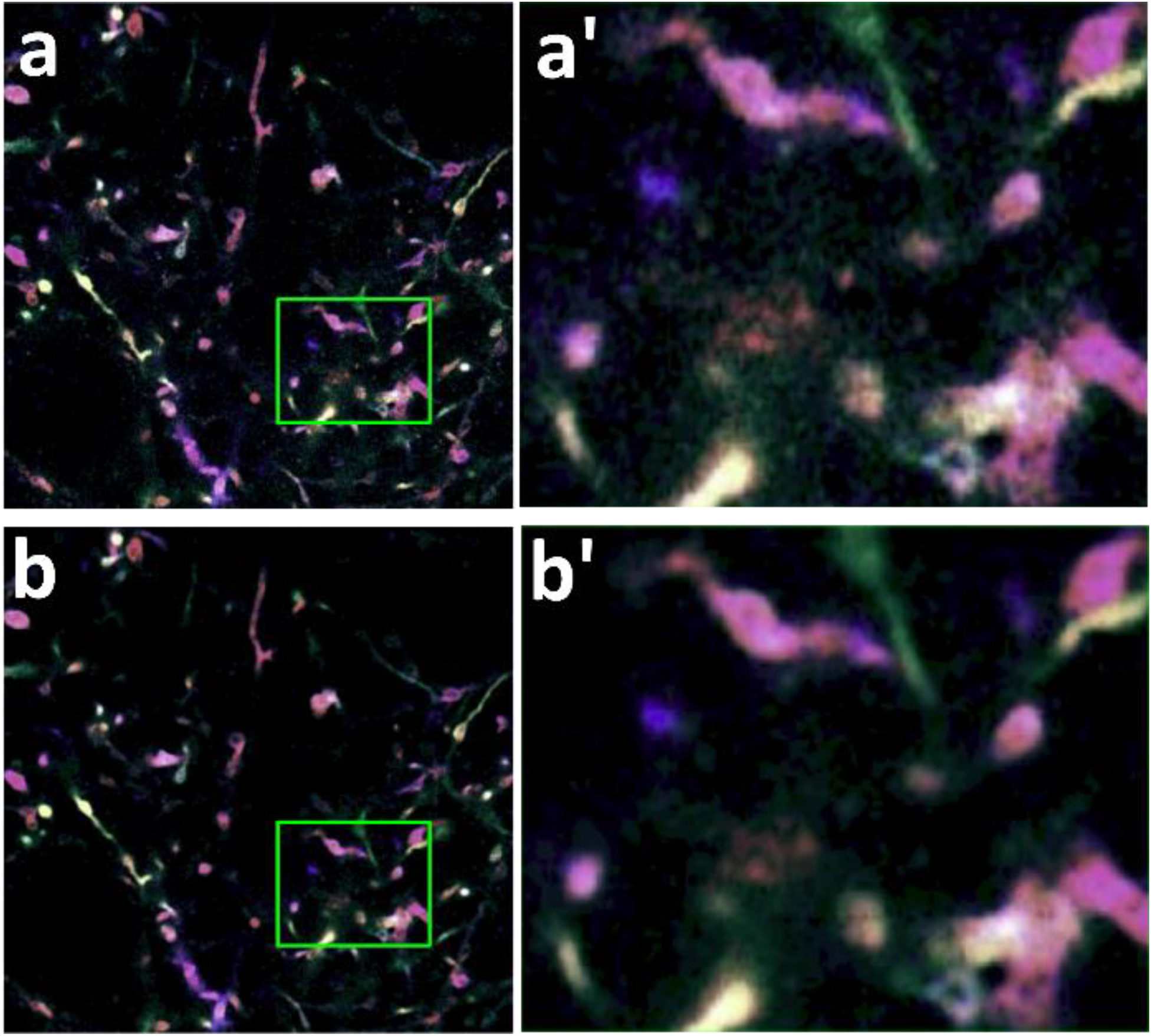
VBM4D denoising of Brainbow image. (a, b) Maximum projection of three adjacent Brainbow Z-slices and its denoised image by VBM4D, respectively. (a’, b’) Magnified views of the green boxes in (a, b), respectively.

### Tri-layer LSTM

For general-purpose sequence modeling, LSTM as a special RNN structure has been proven stable and powerful for modeling long-range dependencies (Hochreiter and Schmidhuber 1997; Sutskever, Vinyals, and Le 2014). LSTM utilizes a memory cell *c_t_* as an accumulator of the state information that is accessed, written and cleared by several self-parameterized controlling gates. Every time a new input comes, its information will be accumulated to the cell if the input gate *i_t_* is activated. Also, the past cell status *c*_*t*−1_ could be forgotten in this process if the forget gate *f_t_* is on. Whether the latest cell output *c_t_* will be propagated to the final state *h_t_* is further controlled by the output gate *o_t_*. One advantage of using the memory cell and gates to control information flow is that the gradient will be trapped in the cell (also known as constant error carousels [24]) and be prevented from vanishing too quickly, which is a critical problem for the vanilla RNN model.

The key equations are shown below:

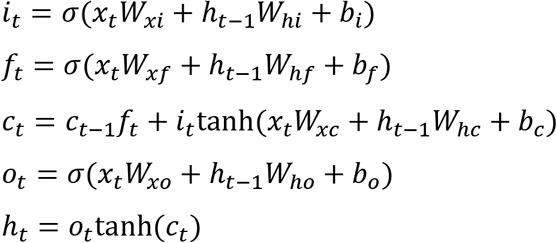

where *x_t_* is the input to the memory cell layer at time *t. i_t_* is input gate at time *t. f_t_* is the activation of the memory cell’s forget gate at time *t. c_t_* is the candidate value for the states of the memory cell at time *t. o_t_* is the value of output gate. *h_t_* is the final output. *Ws* are weight matrices and *bs* are bias vectors. *σ* and tanh are function transformations.

Multiple LSTMs can be stacked and temporally concatenated to form more complex structures. In this paper, we stacked LSTMs into a tri-layer structure (Fig. 1c). The tri-layer LSTM structure is shown in Fig.1c. In our experiments, the inputs to LSTM are the concatenated pixel values of small cubes along the tracing route. The outputs of LSTM are the center coordinate of these small cubes.

## Results

### Brainbow 3D image dataset and ground tracing truth

By randomly expressing different ratios of fluorescent proteins in different spectra, Brainbow stochastically labels individual neurons with distinct colors (Cai et al. 2013; Livet et al. 2007). Therefore, the intermingled neuronal process in the same brain can be differentiated by their distinct spectral combinations. This sheds light to tracing densely labeled neurons for mapping connectomics, i.e. neural connections in the brain. We collected a mouse Brainbow 3D stack, which contains 136 z-slice images. This dataset contains more than 1,000 neuronal processes in four distinct spectral channels. To acquire tracing ground truth, multiple tracers manually reconstructed and validated the soma and neurite in this Brainbow images.

### Implementation details

Our framework was implemented in Keras (Chollet 2015). Keras is a high-level deep neural networks API, written in Python for running on top of either TensorFlow (Abadi et al. 2016) or Theano (Theano Development Team 2016). We use its Tensorflow back-end in our implementation. Experiments were conducted on a workstation with 12G NVIDIA GeForce GTX TITAN X GPU. About half of the ground truth tracing results were used for training and the rest were used for testing.

### Tracing outcome

Automated neural tracing performance can be evaluated quantitatively by using the DIADEM metric (Gillette, Brown, and Ascoli 2011). The DIADEM metric compares, on the basis of topology, two digital morphological reconstructions of one and the same neuron. The two reconstructions are both spatially registered to the same image. stack and thus to each other. The metric is a multi-step process that scores the connection between each node in the gold standard (manual tracing) reconstruction based on whether or not the test (automated tracing) reconstruction captures that connection. Table 1 shows the results computed as the average and standard deviation of DIADEM metric from the ground truth (gold) results to our tracing results (Gold to Predict), and from our tracing results to the gold (Predict to Gold) results. The imaging noise and faint labeling level of certain neurites lead to false positives (i.e., false traced neurites) and false negatives (i.e., missed neurites), which caused most of the tracing error.

**Table 1.**
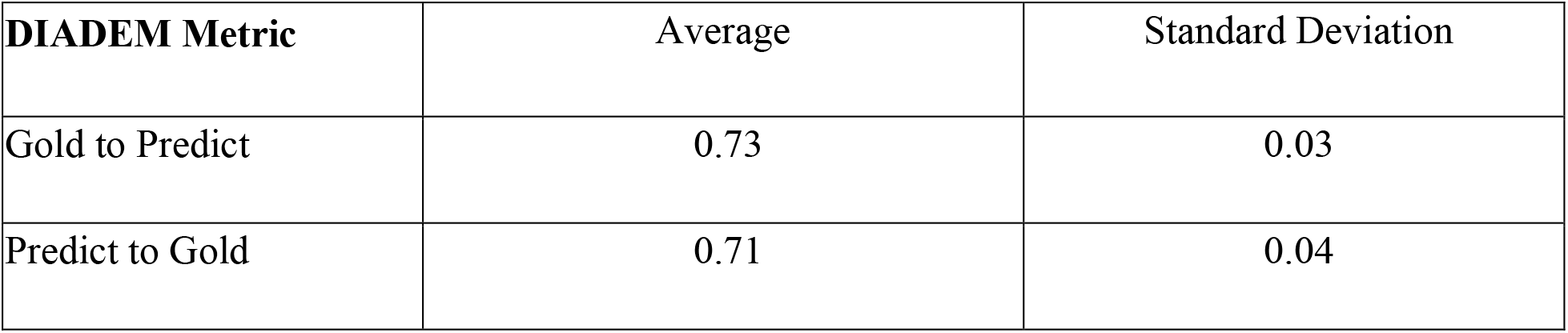
Tracing with gold results using DIADEM metric.

The LSTM gates recursively feed forward their hidden state and the next window value. The final gate passes the hidden state to a dense layer that outputs the prediction. The window size and the hidden layer size are hyperparameters along with optimization parameters such as the learning rate and number of iterations. We evaluated the distance errors with different hyperparameters as shown in Fig.3. Different LSTM window size and hidden layer size will affect the tracing performance. We observed that the best performance is achieved when the window size is equal to 3 and hidden layer size is equal to 128. Too large or too small LSTM window and hidden layer size will hamper the performance.

**Figure 3.**
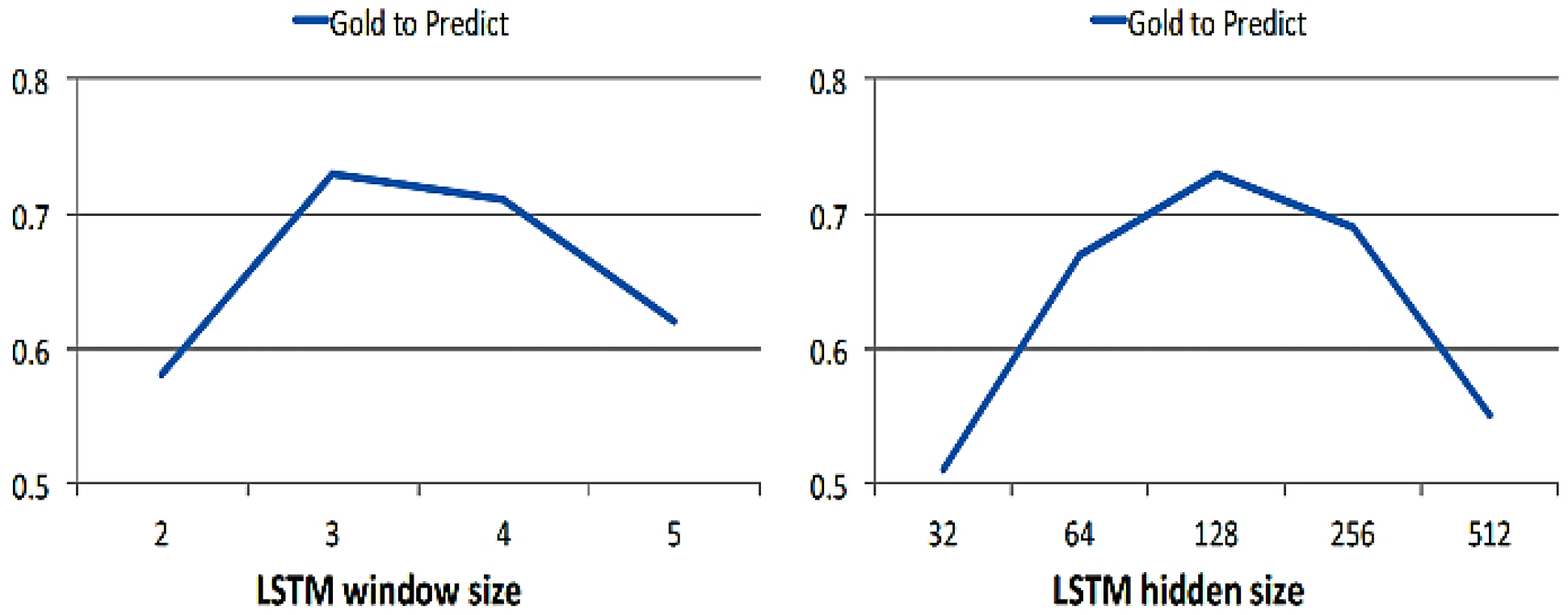
Tracing results for different LSTM window size (left) and different LSTM hidden layer size (right).

Qualitative results are shown in Fig.4. We can observe that our method is able to trace some big soma well. However, there are still some fine-grained important neural parts are missing in the tracing results. This is probably because the information contained in the big soma are more discriminative than normal neurites.

**Figure 4.**
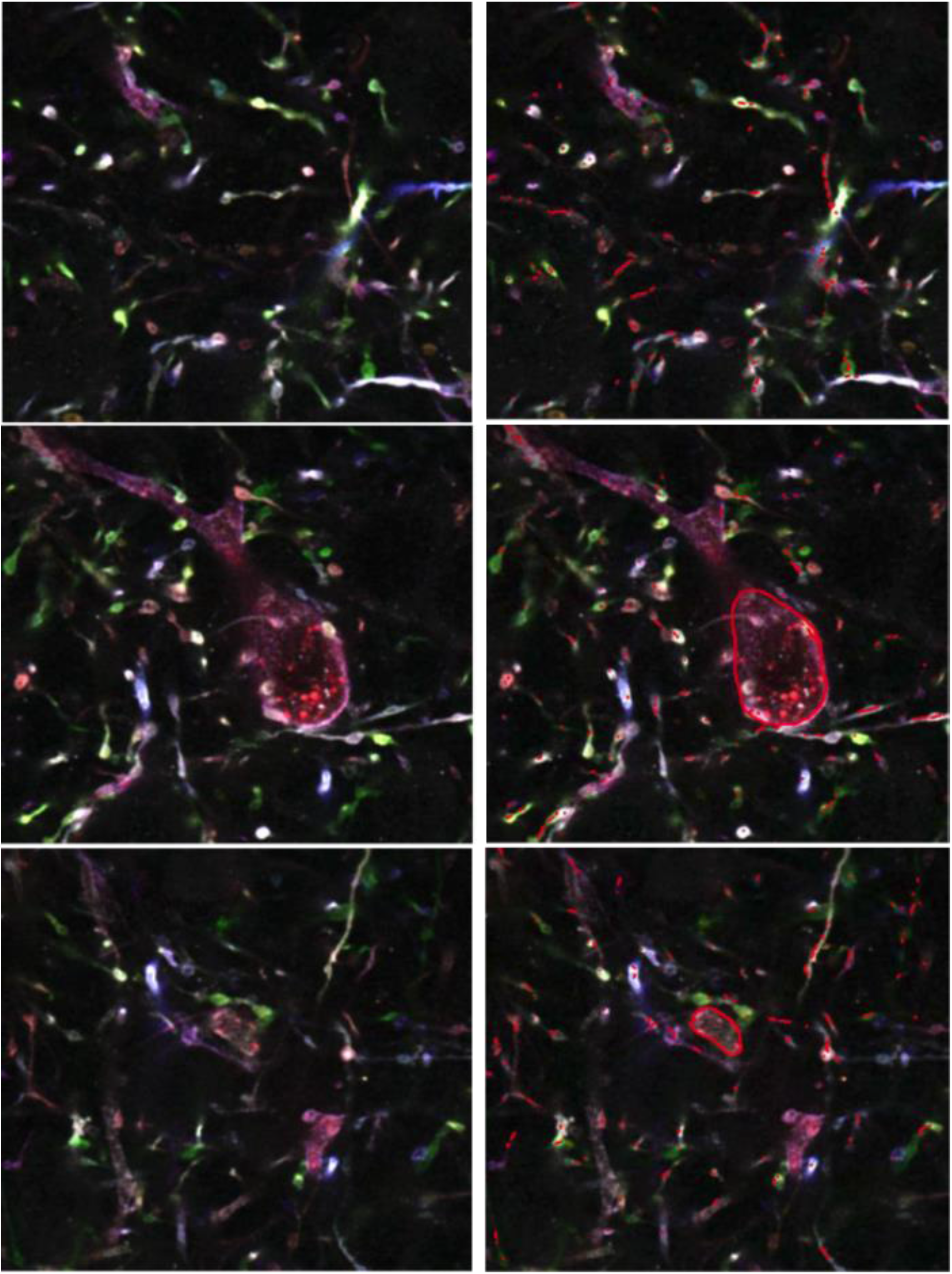
Qualitative results for neural tracing. Original Brainbow slices are shown in the left column. Tracing results (red lines) superimposed on the original slices are shown in the right column.

## Discussion

Neural tracing in fluorescence microscopy images is an important yet challenging problem in neuroscience. Manual tracing of neurons is time consuming and subjective. In this paper, we proposed a recurrent neural network approach for automated neural tracing in Brainbow image stacks. The method contains an image denoising step and a LSTM learning step. The proposed approach is simple and effective. To evaluate our approach, we collected a Brainbow 3D image stack and manually traced a subset to create the gold standard ground truth. We used part of ground truth as training and the rest as testing. Extensive experimental results on the collected dataset demonstrate that the proposed approach performs well for neural tracing in multispectral images. Our framework provides a new paradigm to perform neural tracing based on the merit of LSTM. Currently, we split the denoising step and LSTM tracing learning step. Future work will include the integrating of two steps with a single end-to-end RNN framework.

### Software

3D Brainbow tiff image dataset, tracing ground truth and Python implementation are available at https://www.cai-lab.org/ automated-tracing-lstm.

## Acknowledgements

Y.Y., J.J.C and D.C. were supported by Michigan miBRAIN initiative. D.H.R and D.C. were supported by NIH/NIAI (R01AI130303) and NSF/Neuronex-MINT (NSF-1707316). D.C. was supported by NIH/NIMH (R01MH110932).

